# Omics-analyses of Fermented Onion pickle in Shaping Gut Microbiota and Immune Response in Women: A Community-Based Trial in Pakistan

**DOI:** 10.64898/2026.03.12.711246

**Authors:** Sumbal H. Hafeez, Saad Farooq, Junaid Iqbal, Kumail Ahmed, Sheraz Ahmed, Fayaz Umrani, Sadaf Jakhro, Khaliq Qureshi, Sean R. Moore, Syed Asad Ali, Najeeha Talat Iqbal

## Abstract

A fermented-food intervention trial conducted in Pakistan suggested beneficial changes in the composition of the gut microbiota in healthy women. Using a subset (n=17) of the same participants, this study further investigates the impact of fermented food (onion pickle) on gene expression levels using RNA transcriptomics, with a focus on host–microbiome interactions. After consuming pickles (50g/day) for eight weeks, blood and stool samples of participants were collected at baseline and post-intervention to assess inflammatory markers, 16S rRNA gene sequencing, clinical parameters, and RNA sequencing. Among inflammatory biomarkers, lipocalin (LCN-2) levels significantly decreased (pre=86.5±80.1ng/mL, post=61.0±59.0 ng/mL, p=0.04, paired T-test). Additionally, the intervention downregulated pathways (p<0.05) involved host responses to microbial stimuli, including response to bacterial origin, chemotaxis, and response to lipopolysaccharide. In gut microbiota, observed α-diversity significantly increased post-intervention (p=0.02). Linear discriminant analysis effect size (LEfSe) revealed differential expressions (LDA ≥ 2.0) of *Olsenella* and *Coriobacteriales* at week-8, where *Olsenella sp*. showed a significant negative correlation with LCN-2 (R=-0.36, p<0.05, Spearman’s correlation). These findings suggest that fermented onion pickle consumption for eight weeks modestly alters gut microbial diversity and composition and is associated with reduced inflammatory markers and altered host immune-related gene expression, potentially improving intestinal health.

**Graphical abstract:** 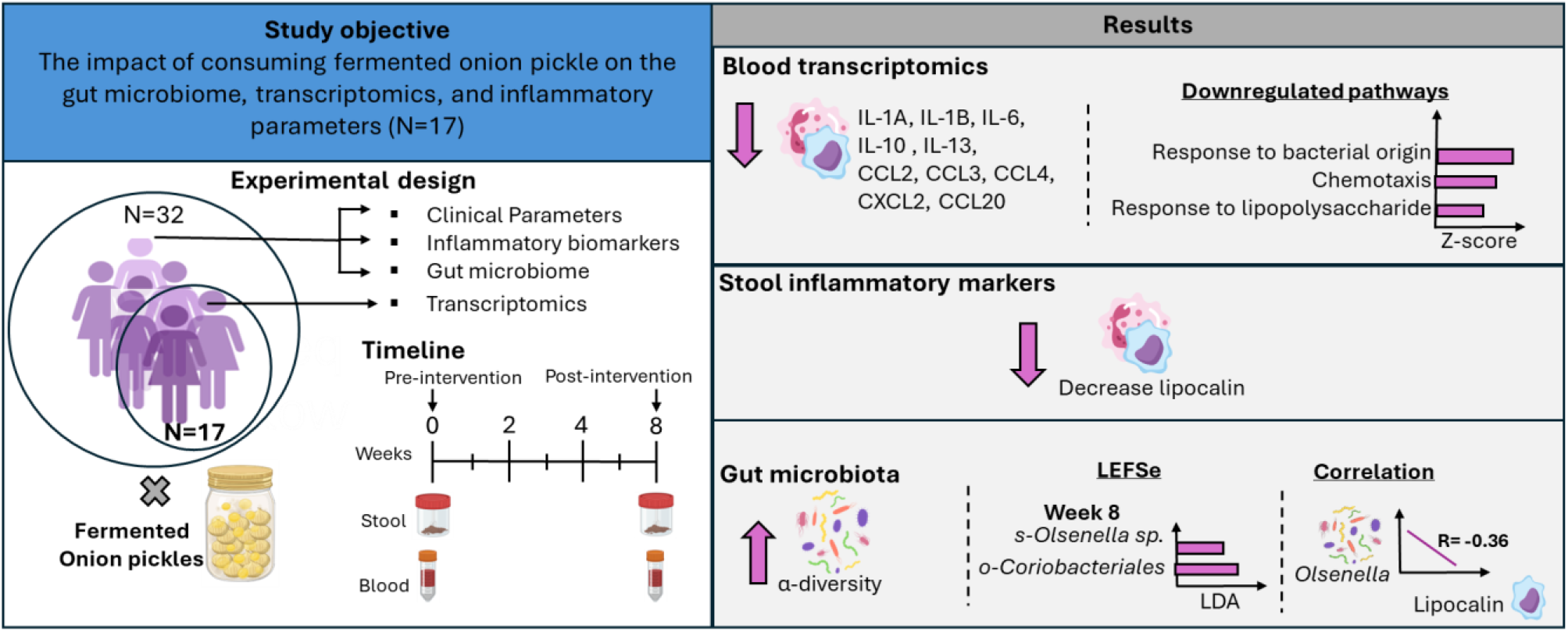

## Introduction

Fermented food is a microbiota-targeted diet, known for its health-promoting properties, as it provides both prebiotics and probiotics to the host gut (1). These components support gut microbial diversity and the immune balance of the host (2). Plant-based fermented foods are valuable for offering low-cost, highly nutritious, and long-shelf-life functional foods, especially beneficial in regions with high malnutrition, food insecurity, and limited access to refrigeration and electricity (1, 3). A variety of fermented vegetable pickles are popular worldwide, particularly in Southeast and South Asia, such as Kimchi in Korea, Paocai (Chinese pickle) in China, Gundruk in Nepal, and various fruit and vegetable pickles known as Achar in India, Bangladesh, and Pakistan (4, 5). Onion, scientifically named Allium cepa, is a rich source of beneficial bioactive compounds like inulin, polyphenols, flavonoids, and carbohydrates, considered as prebiotics (6). The fermentation process not only adds potential probiotics but also enhances the nutritional value of food by inducing physicochemical changes in active constituents, enhancing bioavailability, and reducing FODMAP (Fermentable Oligosaccharides, Disaccharides, Monosaccharides, and Polyols) content (2, 7-9). A study revealed that fermenting onions significantly improved their antioxidant activity, offering various health benefits (10).

Fermented vegetables are a great source of probiotics such as *Lactiplantibacillus pentosus, Lactiplantibacillus brevis, Leuconostoc mesenteroides*, and *Lactobacillus plantarum*, from the genera *Lactobacillus* or *Bifidobacteria* (11). Probiotics are living lactic acid-producing bacteria (LAB), mostly gram-positive and anaerobic rods, that have a positive effect on human health (12, 13). These microorganisms interact with the host immune cells through a complex pathway in the intestine. Probiotics modulate innate and adaptive immunity by regulating dendritic cells, macrophages, T cells, and B cells (14). These are not only involved in the activation and maturation of immune cells but also in immune homeostasis by regulating Treg cells and releasing cytokines such as IL-6, IL-10, IL-17, TNF-α, and IFN-γ (15, 16). Hence, it is crucial to understand the impact of gut microbiome-targeting interventions on the immune system. *In-vivo* and *in-vitro* studies have demonstrated that fermented onions improve colonic histology, increase mucus layer integrity, and enhance antioxidant and anti-inflammatory markers compared to fresh onions (17-19).

Despite promising preclinical results, clinical trials of fermented vegetables in human populations, especially in undernourished settings, are limited. Women of reproductive age (WRA) were chosen as a study group because of the high risk of malnutrition in low-and middle-income countries (LMICs) like Pakistan, where improved nutrition in WRA is essential to effectively reduce maternal and child undernutrition (20, 21). This study aimed to fill this gap by examining the effects of a traditional fermented onion pickle intervention in healthy women living in rural Sindh. This 8-week intervention trial assessed changes in gut microbiota composition, inflammatory biomarkers (myeloperoxidase [MPO] and lipocalin [LCN-2]), and the blood transcriptomic profile of these women.

## Materials and Methods

### Study design and site

This study involves a subset of participants enrolled in a larger clinical trial conducted with 223 women of reproductive age (WRA) living in rural Pakistan, examining the effect of consuming fermented vegetable pickle on gut microbiome and immune parameters. The main study included six fermented pickle intervention groups: Mango-water, Mango-oil, Onion, Radish, Lemon-chilli, and Carrot pickles, each with a minimum of 30 participants, and a control group. From the 32 participants enrolled in the onion intervention group, a subset of 17 participants with RNA samples meeting quality criteria were selected for this study. Approval from the Ethical Review Committee of Aga Khan University was obtained (ERC-2022-6595-23253: Grand Challenges Fermented Food - Achars (fermented pickles) in Pakistan). The trial was registered on ClinicalTrials.gov, identifier: NCT06748313. Informed consent was obtained from all recruited participants.

### Eligibility criteria

Participants under 18 or above 49 years of age, with any gastric/major illness history, who frequently consumed pickles, or antibiotics/probiotics within 2 weeks of the interview date were excluded from the study.

### Intervention and sample collection

The intervention group participants were assigned to consume 50g/day of onion pickle for 8 weeks in addition to their regular meals. Blood samples were collected at weeks 0 (pre-intervention) and 8 (end-intervention) in EDTA tubes for inflammatory markers and transcriptomics analyses. Stool samples were collected in week 0 (pre-intervention) and week 8 (post-intervention) for gut microbiome analyses.

### Nucleic-acid extraction

DNA was extracted according to the manufacturer’s protocol, using a Qiagen DNeasy Power Soil Pro Kit (Qiagen, Germany) and quantified using Invitrogen Qubit 1X dsDNA HS assay kit (Thermo Scientific, USA) on a Qubit Fluorometer.

RNA was extracted from blood samples using the Quick-RNA miniprep plus kit (Cat#R1058, Zymo Research) per the manufacturer’s guidelines. The white blood cell pellet was used to extract RNA for transcriptomics. The purity of nucleic acid was assessed using a Nanodrop 2000 spectrophotometer (Thermo Scientific, USA) and stored at -80°C ULT freezer, until further use.

### Transcriptomics analyses

Out of 32, a subset of 17 participants was selected for RNA sequencing based on the good quality of extracted RNA. Pre and post-intervention samples were shipped to BGI (Shenzhen 518081, China) for RNA sequencing using the DNBSEQ platform. The enrichment of mRNA using total RNA was performed by oligo(dT)-attached magnetic beads. After fragmentation and reverse transcription, double-stranded cDNA was constructed using N6 primers. After phosphorylation and ligation with a bubble adapter, the product was PCR amplified using specific primers. The PCR products were denatured to single strands, and then single-stranded circular DNA libraries were generated using a bridged primer. The constructed libraries were quality-checked and sequenced after passing the quality control.

For the analysis of RNA-seq data, the quality of raw data was assessed using FastQC, and reads were aligned to the human genome GRCh38 using STAR v2.7.11 to check alignment statistics (22). All the samples passed these quality metrics and proceeded to the “quasi-mapping” approach utilized by Salmon for transcript-level quantification (23). This data was imported into R using tximeta (24), which automatically identifies and attaches transcriptome metadata. After preprocessing and exploratory data analysis using Principal Component Analysis (PCA) and unsupervised hierarchical clustering, gene expression analysis was performed in R 4.4.3 (R Core Team 2025) using the DESeq2 package (25). Differential expression analysis was performed based on the DESeq2 negative binomial distribution by comparing post-intervention samples to pre-intervention samples, while controlling for the individuals. Genes with the adjusted p-value threshold of 0.05 or lower and a log fold change of 2 or higher were considered significant Differential Expressed Genes (DEGs). DEGs were visualized using the Bioconductor packages ComplexHeatmap (26) and EnhancedVolcano (27). To explore the possible functions of DEGs, Gene ontology (GO) and Kyoto Encyclopedia of Genes and Genomes (KEGG) pathway enrichment analyses were performed. The significantly enriched KEGG and GO terms were selected for further investigation. Functional enrichment analysis was performed using the clusterProfiler package (28) and web-based portal Metascape (29). The enrichplot package (30) was used for visualization of the GSEA result. The string online database (http://string-db.org) was used to analyze protein-protein interactions (PPI) with a minimum interaction score of 0.4. Cytoscape v3.10.3 was used to visualize the gene network (31) and cytoHubba plugin was used to determine the hub genes within the genes network (32).

### Gut-microbiome analyses

This study presents a transcriptomic subset analysis of participants from a previously published trial of fermented pickles, with targeted gut microbiome analyses included to support integrative host–microbe interpretation. The sequencing data quality and analysis details are discussed in the parent paper (33). The microbiome of the subset data was analyzed using R software (v.4.10). α- and β-diversity indices were measured using the Phyloseq and Vegan packages, respectively. α-diversity (Shannon and Observed) was visualized with a bar plot and analyzed using a paired T-test, while β-diversity was visualized with Principal Coordinate Analysis (PCoA) and assessed by PERMANOVA with 999 permutations. The relative abundance of taxonomic composition was visualized using Taxa barplots created with ggplot2 and phyloseq packages. The graph displayed the top 20 most abundant taxa and grouped less abundant taxa as “others.” Relative abundance data were converted to percentages before plotting.

Differential abundance of bacterial genera between pre- and post-intervention points was analyzed using the DESeq2 package, with shrinkage of log_2_ fold changes performed using apeglm. To reduce noise, taxa were filtered to include only those with more than five counts in at least 20% of samples. Although no taxa met the false discovery rate (FDR) threshold for statistical significance (adjusted p < 0.05), we examined taxa based on their log_2_ fold change (log_2_FC) values to highlight taxa with the most substantial expression changes. A bar plot was generated to show the direction and magnitude of these changes, categorizing phyla/families/genera as upregulated (log_2_FC > 0) or downregulated (log_2_FC < 0). This exploratory analysis offers descriptive insights into taxa that may be biologically relevant, even if they did not reach statistical significance. To further identify microbial markers associated with each intervention point, Linear Discriminant Analysis Effect Size (LEfSe) was performed. Markers with significant Kruskal-Wallis and Wilcoxon signed-rank sum test (p ≤0.05), followed by LDA to estimate effect size at log (10) values (cut-off was set to 2.0), were visualized on a bar plot. The cladograms were plotted using the ggdendro package in R. Spearman correlation was conducted between Lefse markers and inflammatory biomarkers, and a heatmap was plotted using the “pheatmap” package in R.

### Statistical analyses

Variables other than sequencing data were analyzed using Stata/SE 17.0 for Windows. Continuous variables were summarized as means with standard deviations (mean ± SD), while categorical variables were reported as frequencies and percentages. A paired t-test was used to compare mean values across two timepoints. The p-value between the onion complete group and subset was calculated by an independent T-test for continuous variables. A p*-*value less than 0.05 was considered statistically significant.

## Results

### Demographics and baseline characteristics of the intervention subset

Seventeen participants from the onion pickle intervention arm were selected for RNA sequencing based on RNA quality. The baseline demographic and anthropometric characteristics of the RNA-seq subset are summarized in ***Table 1 (Table S1)***. The mean age of participants in the subset was 26.3 ± 6.4 years, with a mean BMI of 20.7 ± 2.7 kg/m^2^. No significant differences were observed between the full onion intervention group and the RNA-seq subset with respect to age, body weight, or BMI, indicating that the subset was representative of the overall study population.

**Table 1:** Demographic and Baseline characteristics of the participants enrolled in the study.

|  | Onion group | Onion sub-set | p-value |
| --- | --- | --- | --- |
| **n** | 32 | 17 | - |
| **Age of participant** | $26.7 \pm 5.9$ | $26.3 \pm 6.4$ | |
| **Marital status** |  |  |  |
| **Married** | 27 | 14 | - |
| **Pregnancy status** |  |  |  |
| **Yes** | 4 | 3 | - |
| **SES (Wealth scores)** | $0.1 \pm 1.1$ | $0.2 \pm 1.0$ | 0.72 |
| **Systolic pressure** | $107.1 \pm 7.9$ | $107.7 \pm 7.1$ | 0.77 |
| **Diastolic pressure** | $73.2 \pm 8.5$ | $73.5 \pm 7.6$ | 0.90 |
| **Weight (kg)** | $49.7 \pm 10.0$ | $47.2 \pm 7.1$ | 0.31 |
| **BMI (kg/m<sup>2</sup>)** | $21.7 \pm 4.1$ | $20.7 \pm 2.7$ | 0.31 |

Data presented as mean ± SD and n (%). \* $p < 0.05$ . SES = Socioeconomic Status.

### Clinical Parameters and Inflammatory Biomarkers

The blood analyses of the participants showed no significant difference in the clinical parameters post-intervention. The analyses of stool inflammatory biomarkers had no significant change in MPO; however, LCN-2 decreased significantly from 86.5 ng/mL to 61 ng/mL (*p=0*.*04*), post-intervention (***Table 2 and Figure S1a***).

**Table 2:**
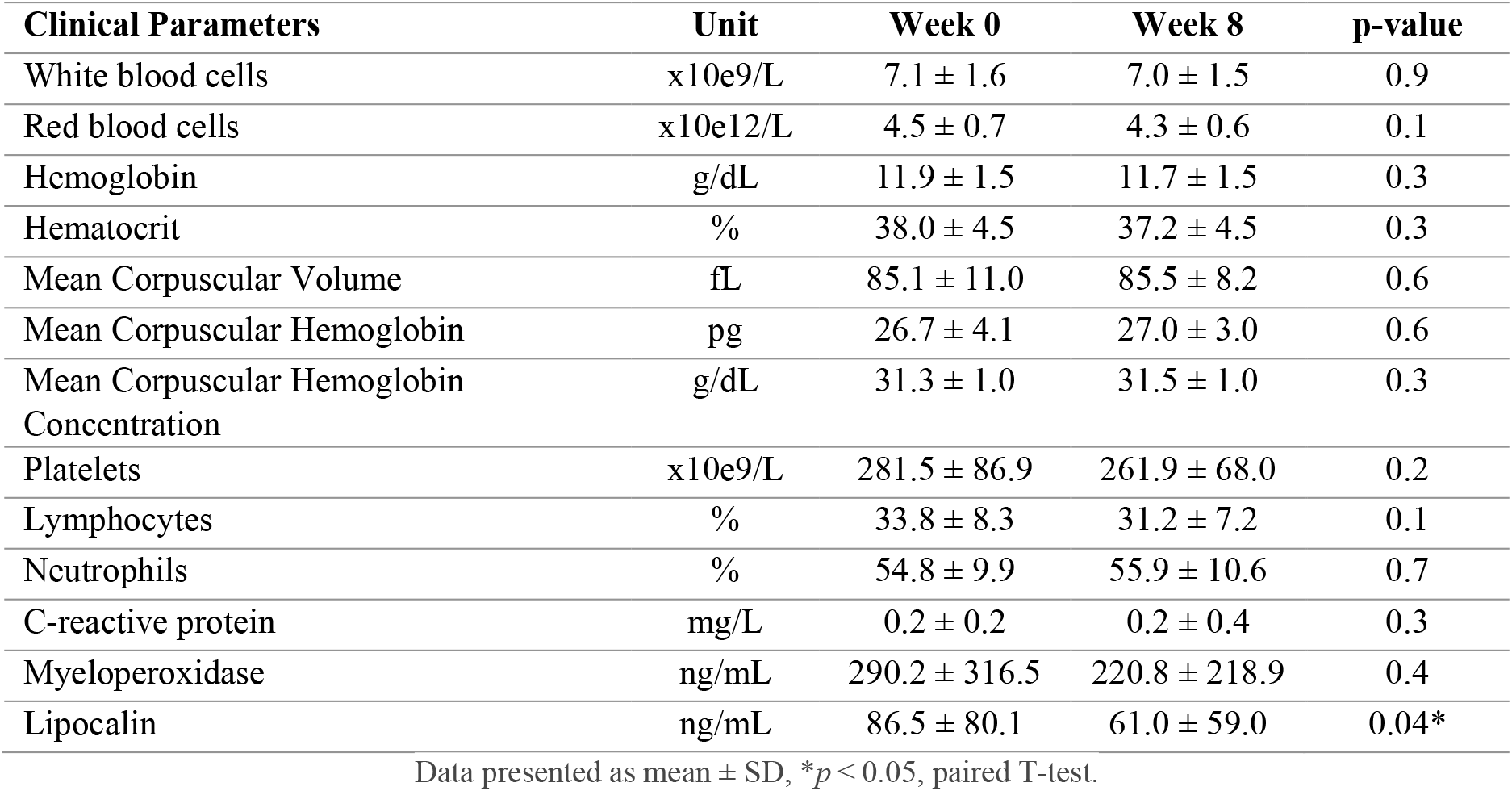
Pre- and post-intervention clinical parameters of the participants.

| Clinical Parameters | Unit | Week 0 | Week 8 | p-value |
| --- | --- | --- | --- | --- |
| White blood cells | x10e9/L | 7.1 ± 1.6 | 7.0 ± 1.5 | 0.9 |
| Red blood cells | x10e12/L | 4.5 ± 0.7 | 4.3 ± 0.6 | 0.1 |
| Hemoglobin | g/dL | 11.9 ± 1.5 | 11.7 ± 1.5 | 0.3 |
| Hematocrit | % | 38.0 ± 4.5 | 37.2 ± 4.5 | 0.3 |
| Mean Corpuscular Volume | fL | 85.1 ± 11.0 | 85.5 ± 8.2 | 0.6 |
| Mean Corpuscular Hemoglobin | pg | 26.7 ± 4.1 | 27.0 ± 3.0 | 0.6 |
| Mean Corpuscular Hemoglobin Concentration | g/dL | 31.3 ± 1.0 | 31.5 ± 1.0 | 0.3 |
| Platelets | x10e9/L | 281.5 ± 86.9 | 261.9 ± 68.0 | 0.2 |
| Lymphocytes | % | 33.8 ± 8.3 | 31.2 ± 7.2 | 0.1 |
| Neutrophils | % | 54.8 ± 9.9 | 55.9 ± 10.6 | 0.7 |
| C-reactive protein | mg/L | 0.2 ± 0.2 | 0.2 ± 0.4 | 0.3 |
| Myeloperoxidase | ng/mL | 290.2 ± 316.5 | 220.8 ± 218.9 | 0.4 |
| Lipocalin | ng/mL | 86.5 ± 80.1 | 61.0 ± 59.0 | 0.04* |

Data presented as mean ± SD, \* $p < 0.05$ , paired T-test.

### Fermented onion pickle consumption modulated the blood transcriptome profile

RNA-seq data from the 17 participants [baseline (n=17) and post-intervention (n=17)] were analyzed to identify host gene expression changes in response to fermented onion pickle consumption. After pre-processing, principal component analysis (PCA) was performed to visualize the separation of pre- and post-intervention samples based on the variation in gene expression among these samples. The plot showed the segregation of pre- and post-intervention samples, where PC1 accounted for 30% of the variance, and PC2 accounted for 8% of the variance **(*Figure 1a*)**.

**Figure 1:**
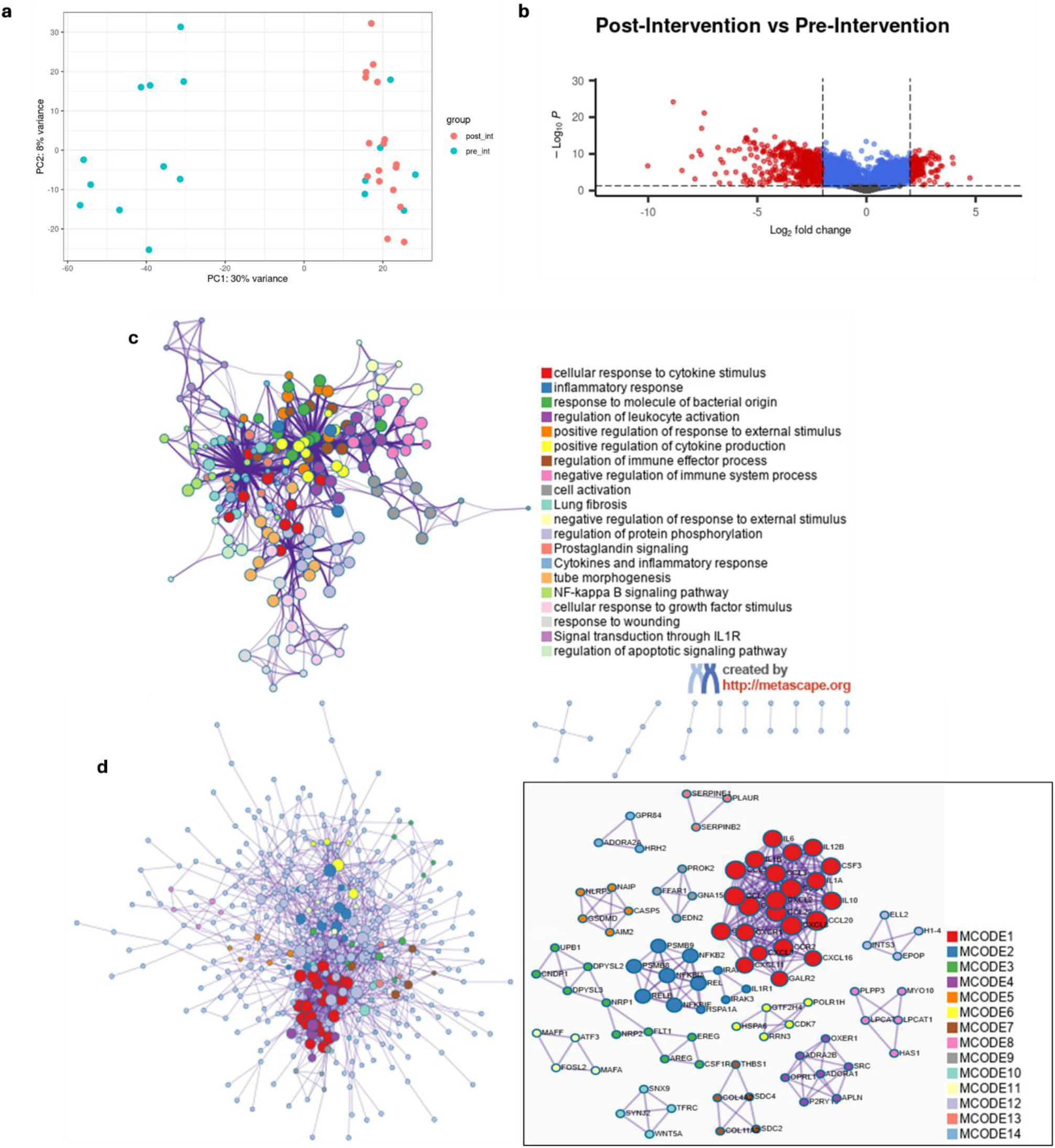
Transcriptomic Overview and DEG Identification: a) Principal component analysis (PCA) showing segregation of pre and post-intervention RNA samples based on the variance in gene expression profile. b) Volcano plot of Differentially expressed genes (DEGs) (Legend: Blue: non-significant, Red: Significant log2FoldChange only, Light blue: Significant p-value only, Red: Significant p-value and log2FoldChange). c) Term enrichment network: each term represented by a node; size proportional to the number of input genes that come under the same term, color represents its cluster identity, and thickness of the edge represents the similarity score. d) Protein–protein interaction (PPI) network of DEGs, highlighting densely connected modules identified using the MCODE algorithm. Gene Ontology enrichment analysis was applied to each MCODE component, retaining the top three terms based on adjusted *p*-values.

Differential gene expression between pre- and post-intervention samples was assessed using DESeq2. Following the removal of lowly expressed features, the dataset for statistical testing consisted of 28,281 genes. Our analysis identified 635 genes that were significant and differentially expressed, with 469 downregulated and 166 upregulated genes in the post-intervention group (***Figure 1b, Table S2***).

To characterize the biological processes affected by the intervention, all DEGs were subjected to over-representation analysis (ORA). Enrichment analysis revealed significant modulation of immune- and inflammation-related biological processes, including cellular response to cytokines, inflammatory response, response to bacterial origin, and leukocyte activation ***(Figure 1c, Table S3)***.

To further investigate the molecular interactions underlying these transcriptional changes, a protein– protein interaction (PPI) enrichment analysis was performed. The network developed clusters that identified proteins that form physical interactions. Densely connected modules were detected using the Molecular Complex Detection (MCODE) algorithm. The three terms with the lowest p-values were retained as the functional descriptions of the network and MCODE components. The highly dense cluster, MCODE1, included cytokine-cytokine interaction pathway (-log10P = 43.1) and Interleukin-10 signaling (-log10P = 41.6) **(*Figure 1d, Table S4*)**. Collectively, these results indicate that fermented onion pickle consumption is associated with transcriptional modulation of immune- and inflammation-related pathways.

Functional analysis identified significantly enriched biological processes associated with the differentially expressed genes (***Figure 2a***). The top biological processes included response to molecule of bacterial origin (gene count 46, (gene count 46, *p*adj = 1.59E-12), response to lipopolysaccharide (gene count 46, *p*adj = 9.08E-12), and leukocyte migration (gene count 43, *p*adj = 9.08E-12) (***Table S5***). Functional enrichment network analysis revealed a dominant immune-related module centered on granulocyte migration, representing modulated innate immune recruiting signals (***Figure 2b***). Consistent with granulocyte migration, the string network analysis revealed that most enriched downregulated genes were chemokine-mediated leukocyte recruitment (CXCL2, CCL20, CCL3, CCL4, CCL2) and inflammation-related genes (IL-6, IL-1A, IL-1B, IL-13, IL-10), suggesting a downregulation of the overall immune response post-intervention (***Figure 2c-2d***).

**Figure 2:**
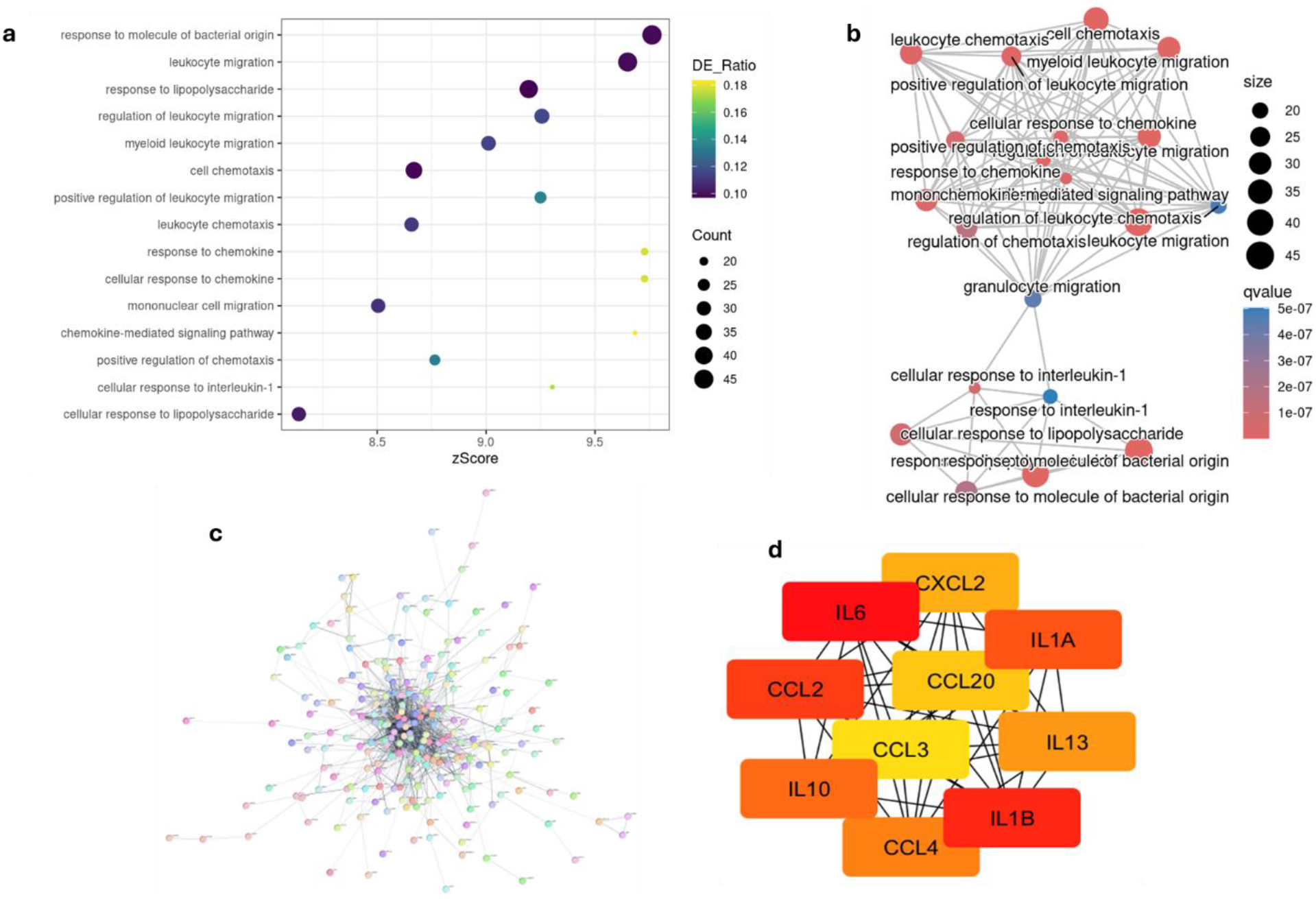
Functional Annotation and Network Analysis of Downregulated Genes: a) Dot plot of significantly enriched biological pathways, with gene set z-scores on the x-axis and pathway terms on the y-axis; dot size represents gene count. (b) Network enrichment analysis illustrating interactions among significantly enriched biological processes. (c) STRING-based interaction network of downregulated genes. (d) STRING network of the top 10 downregulated genes following the intervention.

Downregulated genes were subjected to Gene Ontology (GO) functional analysis to identify biological pathways affected by the intervention. The most downregulated genes were associated with responses to bacterial origin, chemotaxis, and lipopolysaccharides, whereas upregulated genes belonged to IL-1β production and dendritic cells migration (***Figure 3a***). This suggests a coordinated suppression of innate immune activation and cell recruitment pathways, particularly those related to pathogens, and upregulation of pathways involved in tissue repair, contributing to improved immune homeostasis. Gene Set Enrichment Analysis (GSEA) was subsequently performed on the complete ranked gene list to identify pathways affected by the intervention. The cytokine-cytokine receptor interaction (NES=-1.9, *q-value=*6.40E-07), TGF-β (NES=-2.18, *q-value=*6.40E-07), and IL-17 (NES=-2.25, *q-value*=7.43E-07) pathways were identified. These pathways were negatively enriched with genes in the set, concentrated at the lower end of the ranked list, indicating a coordinated downregulation of cytokine-mediated signaling and immunoregulatory mechanisms post-intervention (***Figure 3b-d, Table S6***).

**Figure 3:**
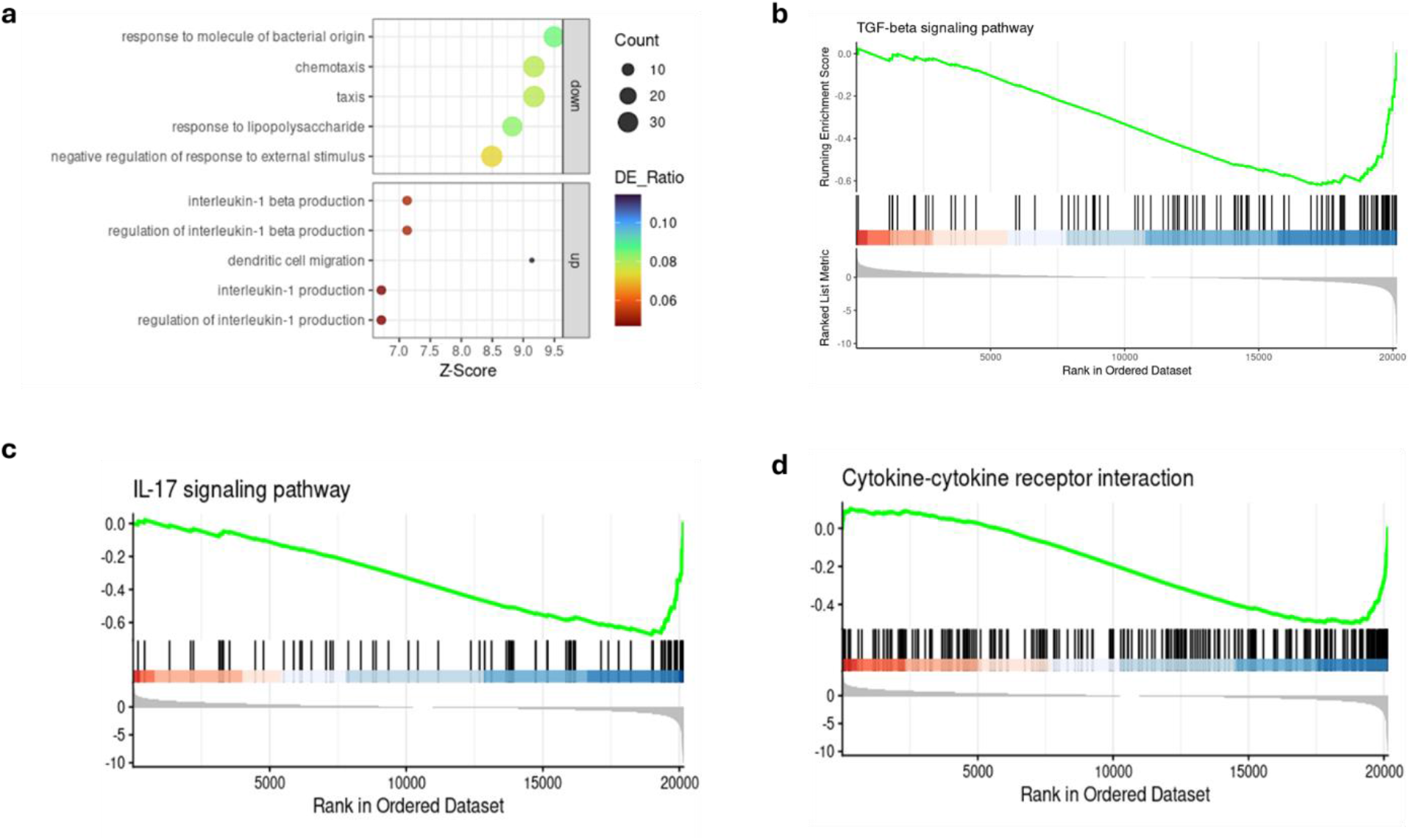
Gene ontology and pathway-level enrichment analysis: (a) Gene Ontology functional enrichment of the top downregulated genes. (b–d) Gene Set Enrichment Analysis (GSEA) plots of the top three pathways significantly affected by the intervention.

### Targeted gut microbiome changes in the transcriptomic subset

To contextualize transcriptomic immune responses, we performed a targeted microbiome analysis in the same subset of participants. The stool DNA of participants collected at baseline (week 0) and post-intervention (week 8) was subjected to 16S rRNA gene sequencing. After quality control, trimming, and merging the dataset using the DADA2 pipeline and assigning ASV using the SILVA database, α- and β-diversity indices were calculated at pre (week 0) and post-intervention (week 8) timepoints. The Shannon α-diversity did not increase significantly in the post-intervention; however, the observed α-diversity index increased significantly (p = 0.02, 95% confidence interval, two-sided, paired T-test) at week 8 compared to baseline **(*Figure 4a, S1b)***. The post-intervention β-diversity was similar to the baseline diversity (***Figure 4b***).

**Figure 4:**
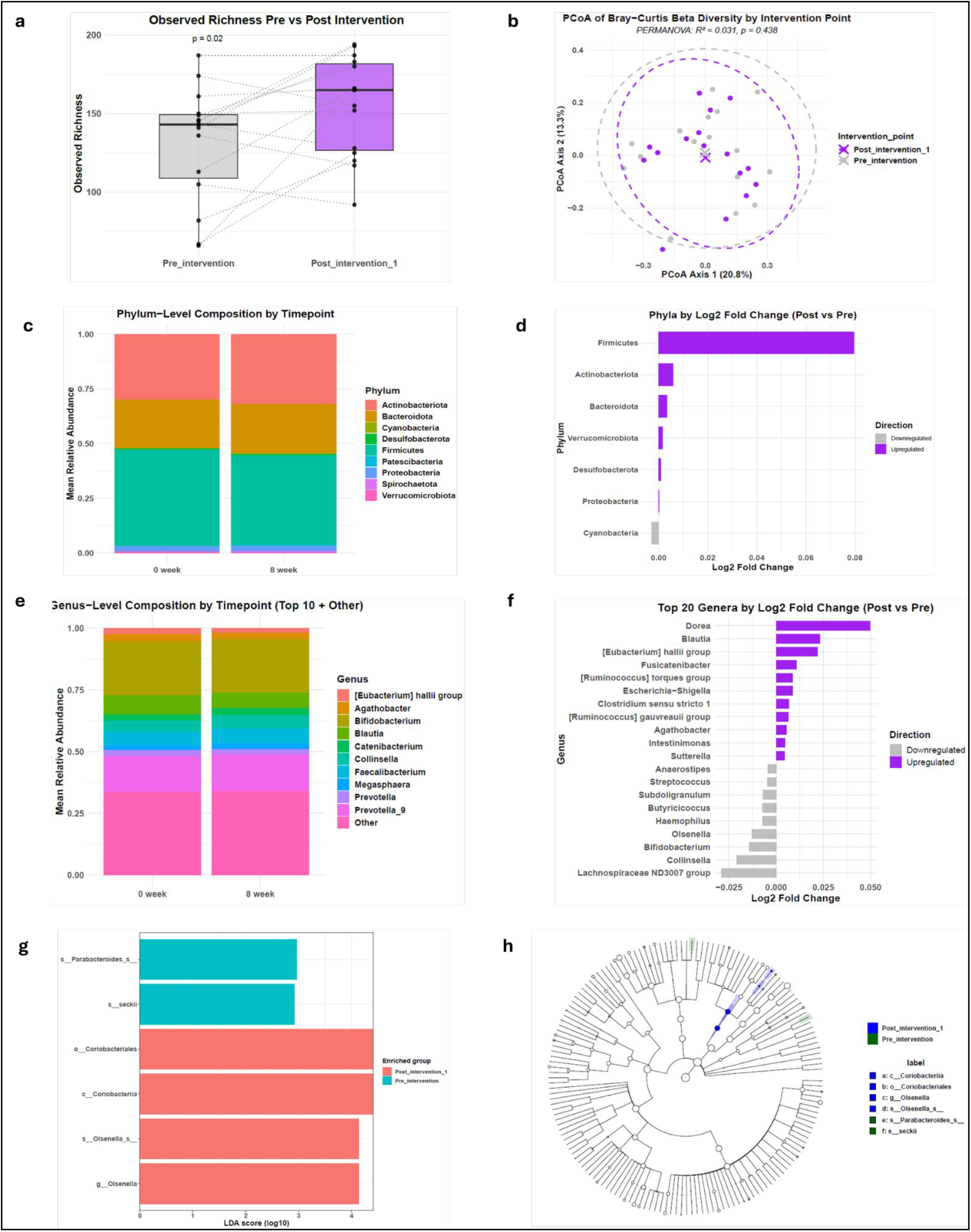
Microbial diversity and compositional analyses: (a) Observed α-diversity (*p= 0.02, n=15 paired, two-sided, paired T-test); (b) β-diversity (p=0.43, PERMANOVA); (c) Relative abundance of phylum-level taxonomic composition at 2 timepoints; (d)) Log_2_ fold-change of the top 20 significantly upregulated and downregulated bacterial families post-intervention; (e) Relative abundance of genus-level taxonomic composition at 2 timepoints; (f) Log_2_ fold-change of the top 20 significantly upregulated and downregulated bacterial genera post-intervention; (g) Linear discriminant analysis size effect (LEFSe) associated with 2 timepoints (Wilkoxon rank test ≤0.05, Kruskal Wallis test ≤0.05, LDA score ≥ 2.0); (h) Cladogram illustrating phylogenetic relationships of LEFSe distinct taxa in the intervention group participants.

To analyze the taxonomic compositional changes over the intervention period, the relative abundance of taxonomic composition was plotted based on the taxa rank count. The taxonomic composition at the phylum level revealed the abundance of phyla *Actinobacteriota, Bacteroidota, Cyanobacteria, Desulfobacterota, Firmicutes, Patescibacteria, Proteobacteria, Spirochaetota*, and *Verrucomicrobiota* at both time points **(*Figure 4c)***. After intervention, there was a slight change in log2fold change of abundance at the phylum level. The relative abundance of phylum *Cynobacteria* decreased, while *Firmicutes, Actinobacteriota, Bacteroidota, Verrucomicrobiota, Desulfobacterota*, and *Proteobacteria* increased **(*Figure 4d)***. Similarly, at family-level, the top 10 families included *Atopobiaceae, Bifidobacteriaceae, Coriobacteriaceae, Eggerthelaceae, Erysipelatoclostridiaceae, Erysipelotrichaceae, Lachnospiraceae, Prevotellaceae, Ruminococcaceae*, and *Veillonellaceae* **(*Figure S1c)***. According to the log2fold change in abundance, Acidaminococcaceae and *Pasturellaceae* increased the most, whereas *Bacteriodaceae* and *Comamonadaceae* decreased **(*Figure S1d)***. The relative abundance at genus-level revealed the top 10 genera, including *Eubacterium (Halii group), Agathobacter, Bifidobacterium, Blautia, Catenibacterium, Collinsella, Fecalibacterium, Megasphaera*, and *Prevotella* **(*Figure 4e)***. According to the log2fold change, Dorea and *Blautia* increased the most, whereas *Lachnospiraceae* and *Collinsella* decreased the most, along with other genera **(*Figure 4f)***.

The Linear discriminant analysis size effect (LEFSe) revealed that post-intervention, the most enriched taxa were *Coriobacteriales and Olsenella*. The pre-intervention differentially abundant taxa included *Parabacteroides* and *Seckii* **(*Figure 4g, Table S7)***. The cladogram showed an abundance of taxa at both time points, along with their hierarchical relationships **(*Figure 4h)***. The inflammatory biomarkers, i.e., LCN-2 and MPO, were correlated with pre and post-intervention LEFSe markers. LCN-2 significantly decreased post-intervention, and Spearman correlation showed that marker 4 (*Olsenella sp*.) was negatively correlated with LCN-2. The unsupervised correlation analysis revealed that all post-intervention markers clustered together, exhibiting a similar correlation profile for biomarkers **(*Figure S1e, Table S8)***. Finally, DEGs were correlated with LEFSe markers, showing non-uniform correlation of markers with genes belonging to NK-cell activation (KIR2DL, KIR3DL families), antigen presentation (HLA-B, HLA-C), and stress-induced immunity (MICA) **(*Figure S1f*)**.

## Discussion

Fermented foods are known to influence gut microbiota and support immune regulation in the host (34, 35). This community trial was conducted in high-malnutrition settings to assess the effect of fermented pickle intervention on the health of women of reproductive age. Previously, this trial presented evidence that regular consumption of 50g of different fermented pickles for eight weeks can significantly remodel gut microbiota, with lemon-chilli and onion pickles being the most beneficial (33). These findings highlight the need to investigate the impact of fermented pickle consumption at the transcriptomics level and to integrate a multi-omics approach. This sub-study further validates our results by demonstrating that regular consumption of onion pickle i) positively affects RNA transcriptome by downregulating immune-activating pathways, ii) improves inflammatory biomarkers profile by decreasing lipocalin, and iii) modulates gut microbiota composition by significantly increasing α-diversity post-intervention.

The primary objective of this study was to investigate the impact of onion pickle intake on the blood transcriptome of women of reproductive age. The immune system is affected by several factors, including individual socioeconomic status, environmental factors, diet, and gut microbiota (36). A study by Zuniga-Chaves et al. found that low SES is associated with less gut microbiome diversity and affinity towards pathologies (37). Therefore, the population living in underprivileged environments often has altered gut microbiota, which leads to immune system activation and low-grade inflammation (38). However, such inflammation can be reduced by improving the gut microbiome through a healthy plant-based diet. A UK study found an association between high intake of vegetables with low lymphocyte profile, 20% of this effect mediated by genus *Collinsella* (39). Similarly, a study by Wastyk et al. compared the effect of a plant-based diet and fermented food diet in healthy participants, and found fermented foods were able to steadily increase gut microbiota diversity and decrease immune markers when compared to a fibre diet (40). Our study hypothesized that consuming fermented onion pickles would lead to a decrease in overall activated immune response, with a specific reduction in pro-inflammatory pathways. Following the intervention trial, our transcriptome results showed significant transcriptional remodelling. Overall, most immune system genes were downregulated (469/635), representing substantial downregulation of the activated immune system. Such transcriptional remodelling is consistent with plant-based dietary studies underscoring the profound influence of nutrition on gene expression (41-43). Our study provided evidence that daily uptake of onion pickle affects host genes by modulating pathways such as cellular response to cytokines, inflammatory response, response to bacterial origin, and leukocyte activation. The enrichment of bacterial response pathways in our study is particularly relevant given the emerging understanding of diet-microbiome-host interactions, where dietary changes alter gut microbiota composition, which in turn influences systemic inflammatory gene expression (44). These pathways are consistent with those identified in another study by Franck et al. following eight weeks of Raspberry consumption (41). The protein-protein interaction (PPI) network analysis identified cytokine-cytokine receptor interaction, inflammatory response, and interleukin-10 signaling as the most significantly enriched functions. The high density of the MCODE1 cluster, which included cytokine-cytokine interaction pathways, proinflammatory and profibrotic mediators, and IL-10 signaling, suggests a coordinated network-level response to the dietary intervention. IL-10, an anti-inflammatory cytokine, plays a critical role in limiting excessive immune responses and maintaining tissue homeostasis (45), contributing to the overall immunosuppressive phenotype observed post-intervention.

Furthermore, the downregulation of key chemokines (CXCL2, CCL20, CCL3, CCL4, CCL2) and interleukins (IL-6, IL-1A, IL-1B, IL-13, IL-10) suggests that dietary intervention engages multiple levels of immune regulation, from chemokine-mediated cell recruitment to cytokine-driven inflammatory amplification. Similar results were observed in participants who consumed fermented foods for 12 weeks, with significant reductions reported in inflammatory interleukins (IL-6, IL-10, IL-18) and chemokines (CCL4, CXCL10, CCL19) (40). The GO pathway analysis of downregulated genes specifically highlighted responses to bacterial origin, chemotaxis, and lipopolysaccharides, indicating a coordinated suppression of pattern recognition and innate immune activation pathways. This is consistent with the hypothesis that dietary intervention modulates gut barrier function and reduces systemic exposure to bacterial products such as lipopolysaccharide (LPS), thereby decreasing the inflammatory burden. These results are in coherence with a study conducted in an animal model where a fruit-vegetable diet was associated with lipopolysaccharide biosynthesis due to a reduction in rgpE-glucosyltransferase protein (46). Lastly, the GSEA identified cytokine-cytokine receptor interaction (NES=-1.9, p.adj=1.05E-06), IL-(NES=-2.25, *p*.*adj=*1.22*E-06*), and TGF-β signaling (NES=-2.18, p.adj=1.05E-06) as negatively enriched pathways. This finding further validates balanced regulation of the immune response, reducing low-grade inflammation, also known as inflammaging, which is associated with various metabolic disorders (47). IL-17 is a pro-inflammatory cytokine (48), whereas TGF-β acts as both a pro- and anti-inflammatory effects depending on the cellular context (49). Simultaneous decrease of these pathways suggests reduced inflammatory stress in the host. Other studies also showed that plant-based diets decreased IL-17, either through PepT1transporter or other mechanisms, indicating that targeted nutritional or microbiota-modulating strategies can attenuate IL-17 signaling in humans (50-52). Additionally, fermented-plant foods are recognized for their immunomodulatory properties, including interactions with TGF-β and Th17 cells mediated by lactic acid bacteria and bioactive compounds, further suggesting that fermentation-derived microbiota shifts can impact host cytokine networks (35).

The second objective of the study was to evaluate the impact of consuming fermented onion pickle on clinical parameters, inflammatory biomarkers, and gut microbiota in the participants. The clinical parameters did not show a significant change following intervention. Similarly, inflammatory biomarkers, including CRP and MPO, did not reach statistical significance; however, LCN-2 was significantly reduced after intervention. A similar effect of CRP was observed in a 6-week clinical trial on women who consumed fermented vegetables (53), and in an 8-week Kimchi (fermented cabbage) intervention trial (54), where circulating CRP levels remained unchanged in human participants. There is no evidence of LCN-2 reduction in fermented food intervention trials, however, a decrease in MPO was found in fermented kimchi and fermented okra intervention trials conducted in animal models (55, 56). The gut-microbiota also showed modulations following the intervention phase. The α-diversity increased significantly, while β-diversity did not show a significant change. An increase in the diversity index was found in contrast with a 4-month clinical trial of vegetable-fruits in healthy adults, where diversity did not change post-intervention (57). However, this finding is in line with fermented foods trial conducted in the Western populations (40, 53). The relative abundance data showed an increase in phylum *Firmicutes*, particularly the genus *Dorea* and *Blautia*, followed by phylum *Actinobacteria*. Both *Dorea* and *Blautia* are short-chain fatty acid (SCFA) producing bacteria, involved in maintaining gut microbiota homeostasis and gut integrity (58). This finding is consistent with a previous trial of fruits and vegetables in US health workers, where *Firmicutes* increased after the intervention (57). Whereas a trial of fermented vegetable intake conducted in healthy German males showed an increase in the phylum *Bacteroidetes* (59). This finding could be linked to the presence of *Lactiplantibacillus pentosus, Lactiplantibacillus plantarum*, and *Pediococcus pentosaceus*, belonging to the phylum *Firmicutes*, in onion pickles offered to our study participants (33). Previous study in Norwegian IBS patients also showed presence of *Lactiplantibacillus plantarum* and *Lactiplantibacillus brevis* in stool samples, when collected after fermented Sauerkraut intervention (60). Lefse analysis revealed enrichment of o*-Coriobacteriales* and g-*Olsenella*, post-intervention. *Olsenella, which* belongs to order *Coriobacteriales* of phylum *Actinobacteria*, is a known SCFA-producing bacteria of the gut (61). A study conducted in a Spanish healthy cohort showed a higher abundance of *Olsenella* in the participants who had adherence to a gut-microbiota targeted diet (62). Correlational analysis between Lefse and inflammatory markers showed a negative correlation between *Olsenella* and LCN-2, suggesting a decrease in gut inflammation by the abundance of *Olsenella* in the gut. There is limited evidence of such association, however, gut bacteria associated with fermentation and SCFA production are inversely related to inflammatory markers, playing key roles in maintaining intestinal homeostasis and modulating host immune responses by suppressing pro-inflammatory signaling and enhancing barrier integrity (63).

This study reveals valuable findings related to the impact of fermented onion intake at the transcriptomic and gut-microbiota level. However, the small sample size and community-specific focus limit the generalizability. Metabolomics, along with RNA seq and microbiome, could have enhanced the mechanistic insights of dietary intervention and its impact on the host. Considering the limited human clinical trial evidence on fermented foods, our study provides valuable insights; however, further mechanistic experiments are needed to elucidate the causal links in host physiology.

## Conclusion

In conclusion, this study demonstrates that fermented onion pickle consumption is associated with coordinated changes in gut microbiota composition and host gene expression, with notable modulation of immune-related pathways. Integrative multi-omics analyses revealed an inverse association between the abundance of the microbial taxon *Olsenella* and the inflammatory biomarker lipocalin, suggesting a link between microbial alterations and reduced inflammatory activity. Although this study was not designed to establish causality, the parallel shifts observed in gut microbiota profiles and host immune gene expression support the presence of a potential microbiota–host immune axis influenced by fermented onion pickle consumption. These observations are further supported by a reduction in fecal lipocalin levels, indicating a concordant attenuation of intestinal inflammatory activity within the same participant subset. Overall, these findings underscore the utility of multi-omics approaches for advancing the understanding of host–microbe interactions in nutritional intervention studies.

## Acknowledgments

The authors are thankful to the study participants for following and completing the study protocol.

## Ethics Statement

Approval from the Ethical Review Committee of Aga Khan University was obtained (ERC-2022-6595-23253: Grand Challenges Fermented Food - Achars (fermented pickles) in Pakistan) (Approval: 02-10-2021). The trial was registered on Clinicaltrials.gov, identifier: NCT06748313 (Registration: 27-12-2024). Informed consent was taken from all the recruited participants. All study procedures were performed according to institutional guidelines and regulations, and following the Declaration of Helsinki.

## Author Contributions

The authors’ responsibilities were as follows: NTI,JI,SA,SAA: designed research; SH,SA,KA,SJ,FU: conducted research; SF,SH: analyzed data; SH: wrote the paper; SAA,NTI: had primary responsibility for final content; KQ: provided data management; SH,SF: provided data interpretation; NTI,SRM,SAA: provided substantial manuscript editing

## Authors Approval

All authors read and approved the final manuscript.

## Data availability

For further data-related information, please contact the corresponding author, NTI, at.

